# Dual indexed design of in-Drop single-cell RNA-seq libraries improves sequencing quality and throughput

**DOI:** 10.1101/835488

**Authors:** Austin N. Southard Smith, Alan J. Simmons, Bob Chen, Angela L. Jones, Marisol A. Ramirez Solano, Paige N. Vega, Cherie’ R. Scurrah, Yue Zhao, Michael J. Brenan, Jiekun Xuan, Ely B. Porter, Xi Chen, Colin J.H. Brenan, Qi Liu, Lauren N.M. Quigley, Ken S. Lau

## Abstract

The increasing demand of single-cell RNA-sequencing (scRNA-seq) experiments, such as the number of experiments and cells queried per experiment, necessitates higher sequencing depth coupled to high data quality. New high-throughput sequencers, such as the Illumina NovaSeq 6000, enables this demand to be filled in a cost-effective manner. However, current scRNA-seq library designs present compatibility challenges with newer sequencing technologies, such as index-hopping, and their ability to generate high quality data has yet to be systematically evaluated. Here, we engineered a new dual-indexed library structure, called TruDrop, on top of the inDrop scRNA-seq platform to solve these compatibility challenges, such that TruDrop libraries and standard Illumina libraries can be sequenced alongside each other on the NovaSeq. We overcame the index-hopping issue, demonstrated significant improvements in base-calling accuracy, and provided an example of multiplexing twenty-four scRNA-seq libraries simultaneously. We showed favorable comparisons in transcriptional diversity of TruDrop compared with prior library structures. Our approach enables cost-effective, high throughput generation of sequencing data with high quality, which should enable more routine use of scRNA-seq technologies.

## Introduction

Most droplet-based single-cell RNA-seq (scRNA-seq) libraries to date have been sequenced on Illumina sequencing platforms using their sequencing-by-synthesis technology (1–4). Libraries generated by droplet-based scRNA-seq approaches require a certain read depth for adequate identification of cell types and states (1–3). With the introduction of Illumina’s NovaSeq6000 next generation sequencing (NGS) platform, the number of scRNA-seq libraries that can theoretically be multiplexed for sequencing together to the required depth has significantly increased (5). Coupled with improvements in hardware technology and sequencing chemistry, sequencing costs can be dramatically reduced, which in turn can facilitate scRNA-seq for routine lab use (Supplementary Table 1). However, the utilization of the improved exclusion amplification (ExAmp) chemistry and patterned flow cells in this new technology has introduced new problems for droplet-based scRNA-seq library structures to date (6–10).

One aspect to be considered when sequencing using ExAmp chemistry is the increased rate of index-hopping between samples sequenced together compared with those sequenced using Illumina’s normal bridge amplification chemistry (7). Index hopping occurs due to the physical incorporation of the sample index from one library into a library molecule from a different library (Fig. 1A–E) (8, 9). The end result is the mis-assignment of reads between samples (Fig. 1B). Index hoppng presents a significant problem for scRNA-seq libraries, where data resolution and sample integrity are vitally important. While computational approaches to use cell barcodes as a second index to solve this mis-assignment problem have been proposed (9, 10), due to the redundant nature of barcodes used in different bead lots, a large amount of data will need to be discarded due to cross-sample barcode collisions detailed below. One of the best strategies to solve the index-hopping problem is to incorporate a second sample index (i5) on the other side of the final sequencing library (Fig. 1F–I) (11). Thus, an index-hopped read would be identified by an un-anticipated combination of sample indexes and can be filtered out. Currently, using a second index and proper sample handling to prevent sample mixing prior to sequencing are the only methods available to pro-actively prevent index-hopping in bulk sequencing assays (8, 11).

**Figure 1.**
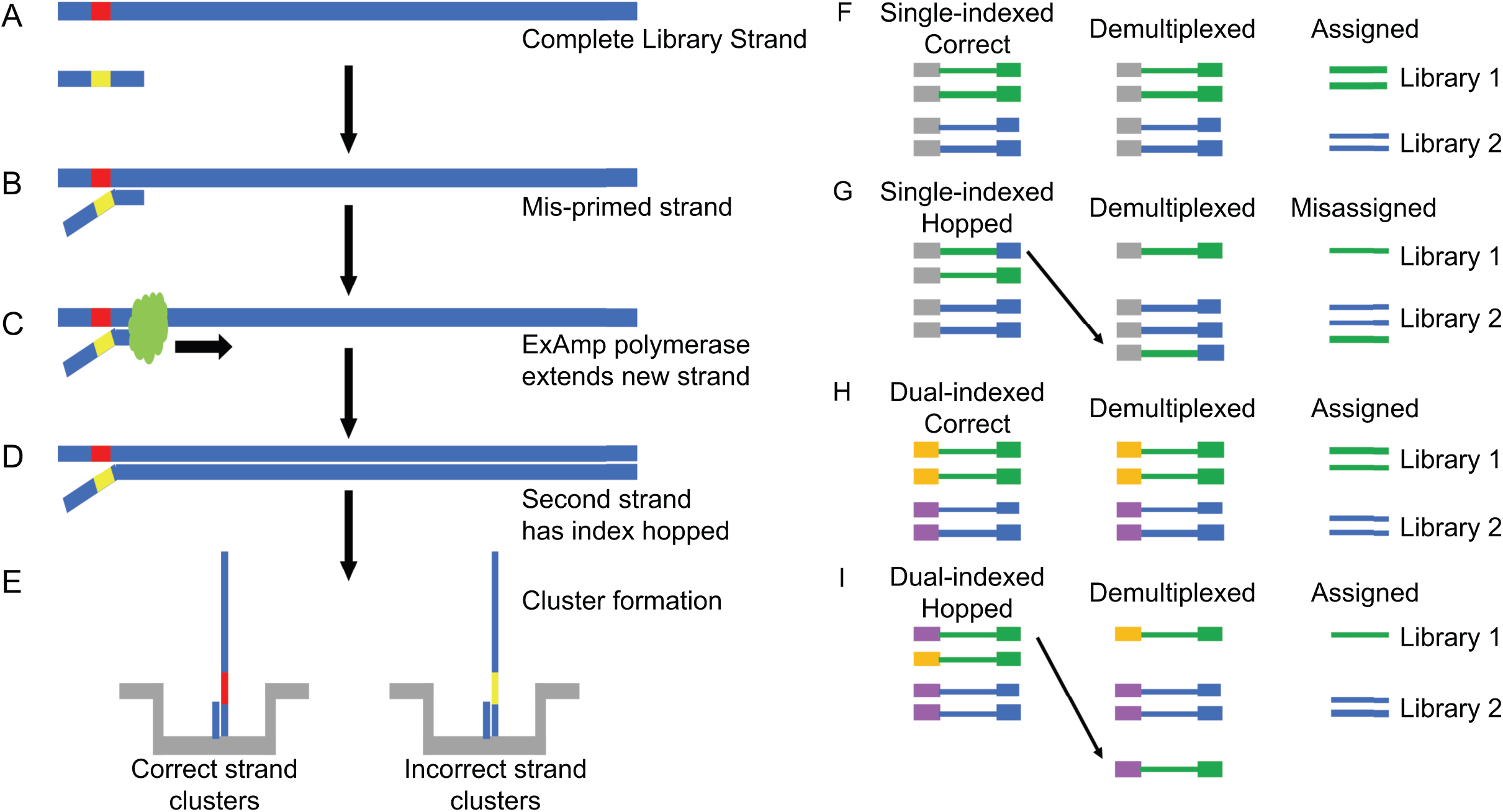
Mechanism for index hopping and its effects on sequencing library demultiplexing. (A-E) Illustration of index hopping due to (A) free adapter molecules remaining after purification post-PCR, resulting in (B) mis-priming of a single stranded library molecule. (C) The mis-primed library molecule is extended via ExAmp polymerase to generate (D) a fully complete library molecule with an incorrect sample index assigned. (E) Both correct and index-hopped molecule can form clusters on the flow cell. (F-I) Demultiplexing runs with single- or dual-indexed libraries with index hopping. (F) The case with a single index and no index hopping where the read(s) for a cluster are associated with a specific sample index (green with green and blue with blue) added to each molecule during library preparation, allowing reads to be assigned to its correct library of origin. (G) The case as above but with index hopping (a blue index now marks a green cluster), where that read will be incorrectly assigned to the wrong library. (H) A unique dual-indexed strategy allows for a single sample to have 2 indexes to be associated with a single library molecule. Here, library 1 = yellow + green, library 2 = purple and blue. (I) The case as above but with index hopping will result in reads displaying unanticipated combination of indexes (e.g., purple + green). The reads associated with unanticipated indexes can then be filtered out.

There are several issues to consider when designing a dual-indexed scRNA-seq library for compatibility with the NovaSeq. A combinatorial dual-indexing scheme in which at least one of the two sample indexes is repeated across two or more samples will reduce the samples that could be potentially mis-assigned. However, samples sharing a sample index would still need to be treated as a single-indexed library (Fig. 1G) (7). The best method then is to use a unique dual-indexed system (Fig. 1I) so that none of the sample indexes on one side of the library (i7) or the other (i5) are shared between samples (7). The indexes used for both sides of the library should be sufficiently different that a 1 base error (insertion, deletion, or substitution) should not result in the mis-assignment of the associated read (12).

Another issue to consider was the use of custom sequencing primers with the prior library structures, such as inDrop V2, that were incompatible with large amounts of other Illumina libraries, such as common TruSeq libraries (2, 13). Thus, previous sequencing runs of V2 scRNA-seq libraries occupy the entire sequencing lanes (Methods). When sequencing just a single library type, the resulting low base composition diversity during the cell barcode read results in a spike in base call error rate. The ability to sequence alongside other Illumina libraries should increase the diversity of bases incorporated across the flow cell at each cycle, improving not only the base calling accuracy, but also the flow cell cluster recognition during sequencing (14).

Here, we document the development and benchmarking of an Illumina compatible dual-index library structure for the inDrop scRNA-seq platform that builds upon the widely-used, commercially available V2 gel beads in a manner independent of the cell barcodes incorporated into the library. We demonstrate the necessity for transitioning to uniquely dual-indexed libraries when sequencing on platforms that use ExAmp chemistry due to cross-sample cell barcode collisions. Using the design documented here, anywhere from 1 to 96 of the resulting scRNA-seq libraries can be sequenced alongside other Illumina samples with minimized sample cross-talk and improvements in sequencing accuracy, which should facilitate the widespread adoption of scRNA-seq in experimental workflows.

## Results

### Sequencing quality of inDrop scRNA-seq libraries is improved when sequenced with a diverse Illumina library

Previously, it was unknown if certain features of inDrop libraries, such as the cell barcodes and spacer region, would interfere with the performance of other Illumina libraries (and vice versa) during sequencing. To assess compatibility with Illumina TruSeq libraries, inDrop V2 libraries were sequenced alongside a 10-15% spike in of Illumina’s PhiX control. Sequencing on both a low-throughput nano run on MiSeq, as well as a mid-throughput NextSeq run, were successful with appreciable number of reads from inDrop V2 libraries (74.6% and 94.2% of the target read depth, respectively; Table 1).

**Table 1.**
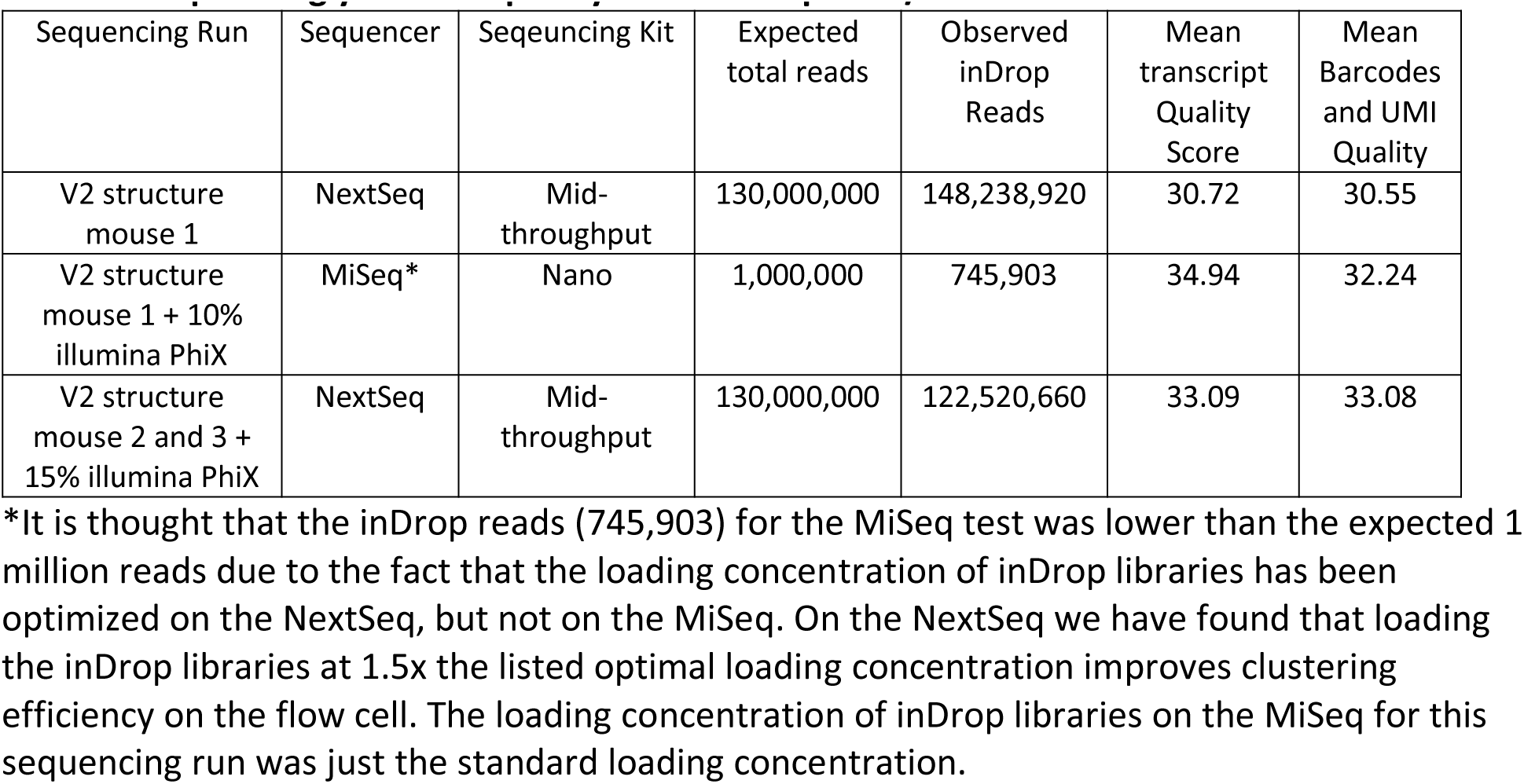
Sequencing yield and quality of V2 inDrop with/without standard illumina libraries

| Sequencing Run | Sequencer | Sequencing Kit | Expected total reads | Observed inDrop Reads | Mean transcript Quality Score | Mean Barcodes and UMI Quality |
| --- | --- | --- | --- | --- | --- | --- |
| V2 structure mouse 1 | NextSeq | Mid-throughput | 130,000,000 | 148,238,920 | 30.72 | 30.55 |
| V2 structure mouse 1 + 10% illumina PhiX | MiSeq* | Nano | 1,000,000 | 745,903 | 34.94 | 32.24 |
| V2 structure mouse 2 and 3 + 15% illumina PhiX | NextSeq | Mid-throughput | 130,000,000 | 122,520,660 | 33.09 | 33.08 |

\*It is thought that the inDrop reads (745,903) for the MiSeq test was lower than the expected 1 million reads due to the fact that the loading concentration of inDrop libraries has been optimized on the NextSeq, but not on the MiSeq. On the NextSeq we have found that loading the inDrop libraries at 1.5x the listed optimal loading concentration improves clustering efficiency on the flow cell. The loading concentration of inDrop libraries on the MiSeq for this sequencing run was just the standard loading concentration.

Importantly, although the PhiX spike-in occupied some of the read depth, the mean quality score increased for the transcript read and barcode + UMI, compared with a run without PhiX (Table 1) (15). The improved quality scores equate to a decrease in the probability of an error in base calling from 8.803 × 10^4^ to 4.917 × 10^4^ on the transcript read, and a corresponding decrease in error probability from 8.455 × 10^4^ to 4.908 × 10^4^ on the barcode + UMI read. This represents about a 1.8- and 1.7-fold decrease in the base calling error rate for bases incorporated during sequencing. This is also reflected in the base calling accuracy plots from the two sequencing runs (Fig. 2). The base calling accuracy plot describes the spread of quality scores as each base is sequenced. It is interpreted as a series of box plots where each box plot maps the percent of clusters in each image of the flow cell with quality scores 2: 30 (called Q30) in each flow cell imaging cycle. When inDrop V2 and Illumina PhiX are sequenced together (Fig. 2B), the transcript read (cycles 1-100) median Q30 barely drops below 80% from cycles 80-100, whereas the inDrop V2 only library median Q30 decrease below 60% during cycles 80-100 (Fig. 2A). In addition, for combined libraries, the Q30 during the barcode + UMI read (cycles 114-164) is maintained at or above 80% for most of the cell barcode + UMI read (Fig. 2B). These results demonstrate that inDrop V2 libraries are compatible with low concentrations of standard Illumina libraries for sequencing and that when sequenced together, the sequencing quality, especially for the non-diverse barcode region, is improved for inDrop libraries.

**Figure 2.**
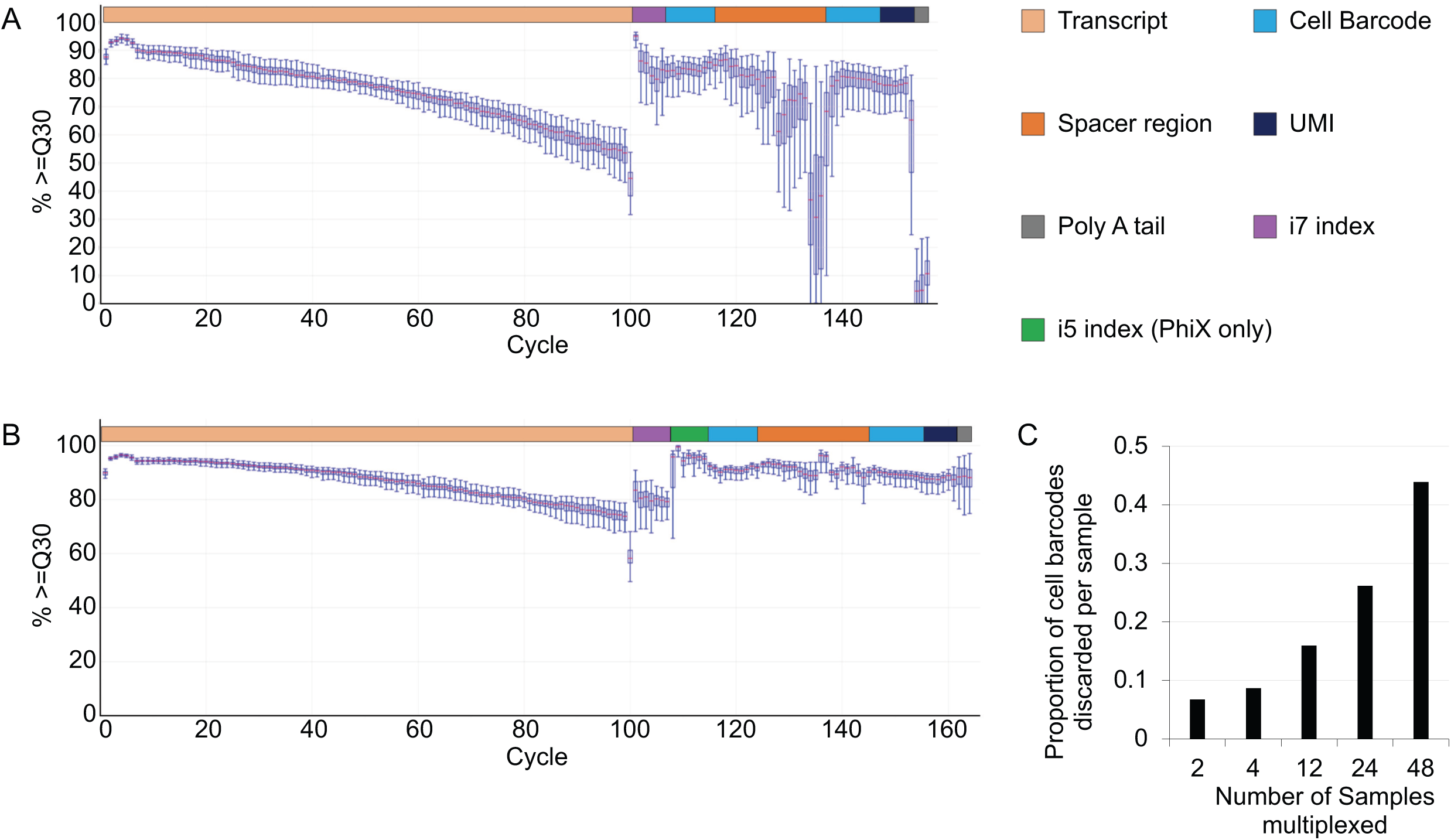
Quality of single-indexed inDrop libraries sequenced alongside Illumina libraries and predicted data loss due to index hopping. (A) The base calling accuracy plot for a V2 inDrop library on a NextSeq sequencing run, depicting the spread of quality scores as each base is sequenced. This plot consists of a series of box plots where each box plot maps the percent of clusters in each image of the flow cell with quality scores ≥ 30 (called Q30) in each cycle. The first 100 cycles correspond to the transcript read; the next 6 correspond to the i7 index read; the final 50 correspond to the cell barcode +UMI reads. The last 6 cycles read into the Poly A tail due to the variable length of the inDrop cell barcodes. (B) The base calling accuracy plot for a V2 inDrop library alongside the control Illumina library, PhiX, on a NextSeq. When sequencing alongside PhiX, the 7-base long i7- and i5-index reads are used so that PhiX reads can be filtered out and discarded during demultiplexing. (C) Plot of the calculated proportion of cell barcodes that will need to be discarded from single-indexed sequencing runs at different levels of multiplexing. We assume each sample will contain ~3000 cell barcodes.

### Redesigned inDrop library structure potentially enables high-throughput NGS

Having demonstrated the compatibility of inDrop libraries with standard Illumina libraries in NGS, we next sought to re-engineer the inDrop library structure for the exclusion amplification (ExAmp) chemistry-based sequencers, such as the NovaSeq6000. Specifically, we sought to incorporate dual-indexing to overcome the well-documented indexing hopping problem on the NovaSeq (3, 6). If two single-indexed samples share cell barcodes and index hopping occurs, then it will be impossible to determine the origins of a particular read belonging to the shared barcode, resulting in the discarding of cells with shared barcodes across indices. We call this problem cross-sample barcode collision, and calculated the theoretical amount of data discarded upon multiplexed NovaSeq runs (Supplementary File 2). For pools of 2, 4, 12, 24, and 48 samples the percentage of cell barcodes, and hence cells, discarded due to cross sample barcode collisions would be 8.67%, 15.99%, 26.19%, and 43.87% respectively (Fig. 2C) (1, 2, 16, 17).

To minimize the possibility of cross-sample barcode collision a second i5 index was incorporated when designing the new library structure. The i5 and i7 indexes used follow a unique-dual indexing strategy such that when only considering one side of the library, each index is only used once. Because part of the Illumina TruSeq read 1 sequencing primer site is built into the oligo used on the barcoded inDrop capture beads (2, 13), it was decided that the newer libraries would use the dual indexed, Illumina TruSeq library structure. The new libraries incorporation of standard Illumina TruSeq adapter sequences (Fig. 3) (13), includes the P5 and P7 flow cell binding sites, the TruSeq standard sequencing primer binding sites (in contrast to prior V2 libraries which require custom sequencing primers), and unique dual indexes (Fig. 3). Furthermore, to achieve a standard Illumina TruSeq library structure, the cell barcode + UMI read has been swapped to read 1, which has been documented to be the higher quality read (18). Since these indexes were designed to be pooled in sets of 8 index pairs (19) and the maximum number of libraries that can be sequenced to a read depth of ~100 million reads per sample on a single lane of the NovaSeq is 25 (5), we selected 24 index pairs to be used as the new indexes in the new library structure. Theoretically the number of usable index pairs could be increased up to 3840 using IDT’s set of 10 bp unique dual indexes, although they would have to be individually validated. We call this new library structure TruSeq-inDrop (TruDrop). The final sequence for the barcode + UMI and transcript sides of TruDrop libraries are as follows: Cell Barcodes: 5’ – AATGATACGGCGACCACCGAGATCTACAC[i5]ACACTCTTTCCCTACACGACGCTCTTCCGATCT[cell barcode 1]GAGTGATTGCTTGTGACGCCTT[cell barcode 2][UMI]TTTTTTTTTTTTTTTTTTT… – 3’. Transcript: 5’ – CAAGCAGAAGACGGCATACGAGAT[i7]GTGACTGGAGTTCAGACGTGTGCTCTTCCGATCTNNNNNN… – 3’.

**Figure 3.**
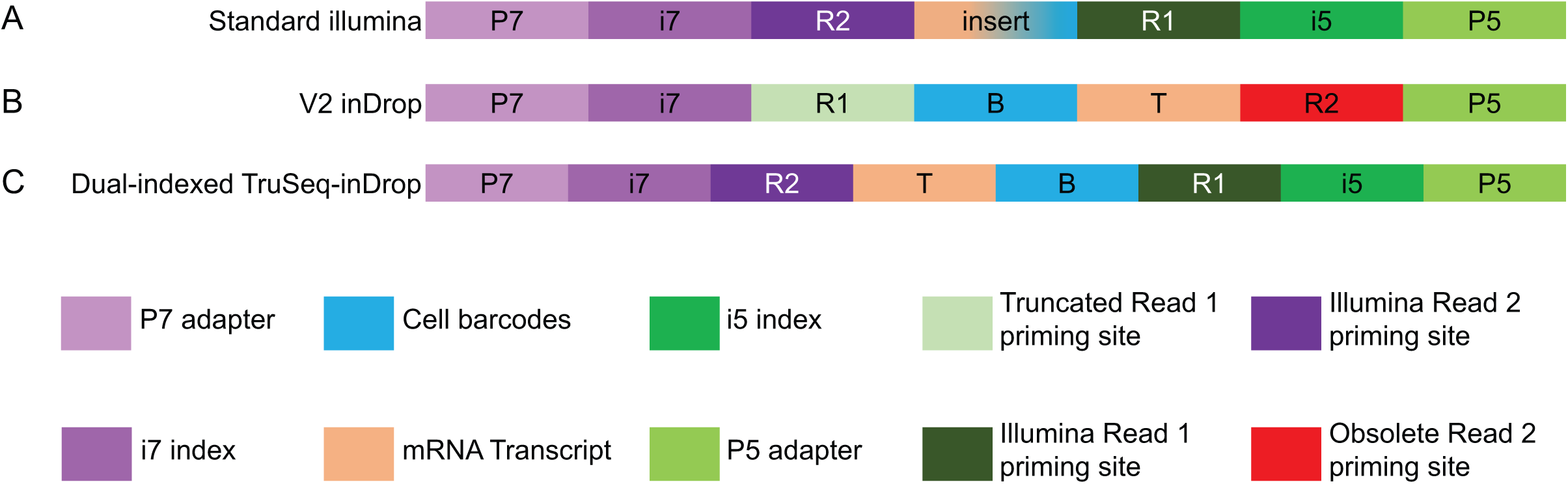
Variations of inDrop library structures from the perspective of sequencing. (A) A standard Illumina library contains P7 and P5 adapter sites that are used to bind Illumina sequencing flow cells. i7-and i5-indexes are incorporated onto the P7 and P5 sides, respectively, to adopt a dual-indexing strategy. On either side of the insert are sites (R1 and R2) where standard Illumina sequencing primers are used to read across both sides of the insert. The reverse complement of these read priming sites then allows for the priming and subsequent reading of the i7 and i5 sample indexes. (B) The V2 inDrop library structure also incorporates the P7 and P5 flow cell adapter binding sites, with a single i7 index. The V2 structure utilizes a R1 priming site that is a truncated version of the standard R2 priming site, and a R2 priming site that is a deprecated R2 priming site. In addition, the R1 and R2 of the V2 structure are flipped so that the insert is read backwards from a normal Illumina library. (C) The TruSeq-inDrop (TruDrop) structure incorporates a second (i5) index and the standard Illumina R1 and R2 priming sites that are used in all Illumina TruSeq libraries.

A detailed version of the custom primers for library preparation, indexes, and methods, and library pooling guidelines used for TruDrop libraries can be found in the supplementary materials.

### TruDrop primers function similarly to V2 primers during inDrop library preparation

As TruDrop uses redesigned primers to generate libraries compatible with TruSeq libraries, it was important to verify that all indexes could be appropriately used to complete and amplify inDrop libraries during the final stages of library preparation. Of the initial 24 tested, all but 1 (TruDrop index pair 9) yielded qPCR amplification curves similar to those of V2 primer pairs (Supplementary Fig. 1A). Furthermore, the Ct values of TruDrop primer pairs 1-8 and 10-24 were well within 1.5 cycles of the average Ct (Supplementary Fig. 1B), suggesting little to no difference in amplification bias between the new primers and the prior V2 primers. As TruDrop index pair 9 failed to amplify in a manner similar to that of libraries with V2 index 6 and 12, it was replaced with index pair 25 (which behaved similar to V2) in all further testing.

### TruDrop libraries see improved performance when sequenced using exAMP chemistry

To put TruDrop libraries into action, we first sequenced these libraries on the iSeq 100, which utilizes patterned flow cells and ExAmp chemistry to test clustering efficiency and priming effectiveness during the sequencing run (20, 21). Two replicates of V2 libraries that had previously performed well on the NextSeq were prepared as TruDrop libraries. The TruDrop samples were then sequenced alongside PhiX on the iSeq 100, yielding an average of 151% of the 2 million reads per library target read depth (Supplementary Table 2). The median Q30 remained at or above 90% during most of the barcode + UMI cycles (cycles 1-11 and 31-50). While for the transcript cycles (cycles 167 – 316), the median Q30 remained at or above 80% for the full 150 cycle transcript read (Fig. 4A). However, if only the first 100 bases of the transcript read (the same length as the NextSeq read length) were considered then 90% or more of reads were above Q30. Thus, it is expected that TruDrop libraries can be sequenced on the NovaSeq but also see improved read quality scores compared with V2 libraries sequenced on the NextSeq with PhiX.

**Figure 4.**
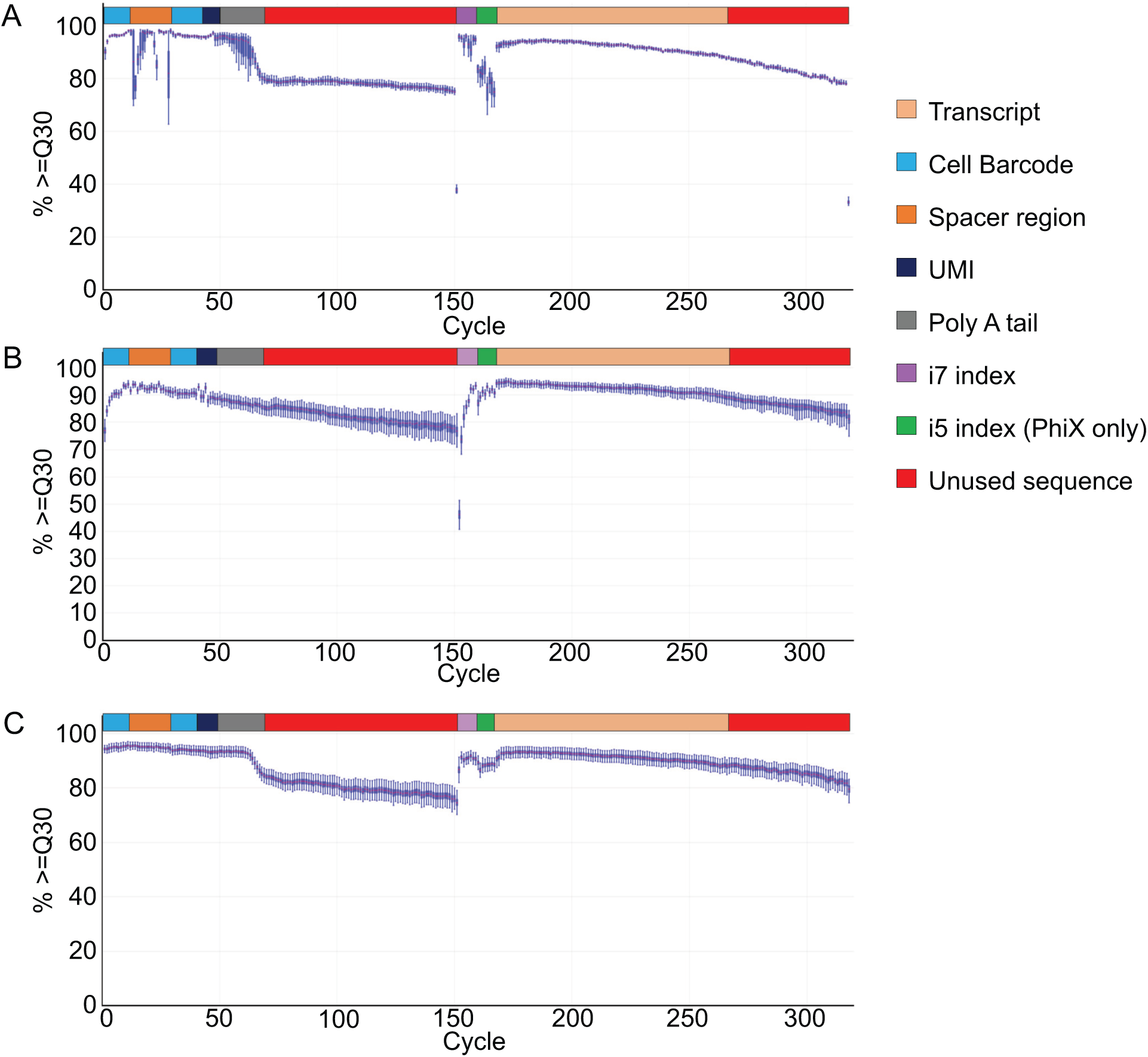
Sequencing quality of TruDrop libraries on exAmp chemistry sequencers. (A) The base calling accuracy plot for two dual-indexed TruDrop libraries on iSeq alongside PhiX. Cycles 1 – 50 depict the quality scores for the cell Barcode + UMI read. Cycles 51 – 151 are sequence data that will be trimmed and discarded during analysis. Cycles 152 – 159 correspond to the i7 index read. Cycles 160 – 167 are the i5 index read. Cycles 168 – 318 are on the transcript read. For the purpose of direct comparison only cycles 168-267 are marked as transcript as only 100 bases of transcript were sequenced for the V2 libraries. (B) The base calling accuracy plot for the same 2 TruDrop libraries when sequenced on the NovaSeq alongside 107 other libraries. (C) The base calling accuracy plot for 24 dual-indexed TruDrop library sequenced on a NovaSeq alongside 186 other libraries.

The same TruDrop libraries were then sequenced on the NovaSeq 6000 alongside 107 other standard Illumina libraries (Table 2). The TruDrop libraries yielded 107% and 89.1%, respectively, of their target read depth (50 million reads), accounting for 0.64% and 0.53%, respectively, of the 3 NovaSeq lanes they were on. Compared to prior tests with V2 libraries on the NextSeq, this was the equivalent of sequencing alongside 99% PhiX with no loss in targeted read depth. In addition, there was an increase of 1.5% – 5.3% in the number of flow cell clusters with perfect index reads compared to V2 libraries on the NextSeq (Table 2). Quality scores were further improved, corresponding to a 2.1- and 1.8-fold reduction in base call error rate compared with sequencing V2 libraries on the NextSeq with PhiX, and a 3.7- and 3.0-fold decrease compared with sequencing just V2 libraries alone on the NextSeq. The base call accuracy plot reflects this improvement (Fig. 4B), as 90% or more of reads that were from TruDrop libraries during read 1 (cell barcode + UMI) and read 2 (transcript) that are of interest in inDrop libraries were at or above Q30. These results demonstrate that not only can TruDrop libraries be sequenced on the NovaSeq, they also see significant improvements in the sequencing quality for both the transcript and barcode + UMI regions.

**Table 2.** Evaluation of 2 TruDrop libraries’ raw yield and quality in sequencing run on the NovaSeq

| Library | TruDrop Mouse 4 | TruDrop Mouse 5 | V2 Mouse 2 + 15% illumina PhiX | V2 Mouse 3 + 15% illumina PhiX |
| --- | --- | --- | --- | --- |
| Sequencer | NovaSeq 6000 | NovaSeq 6000 | NextSeq | NextSeq |
| i7 | CCGCGGTT | TTATAACC | GCCAAT | CTTGTA |
| i5 | AGCGCTAG | GATATCGA | --- | --- |
| Targeted inDrop Read Depth | 50,000,000 | 50,000,000 | 65,000,000 | 65,000,000 |
| Observed inDrop Reads | 53,655,662 | 44,554,464 | 57,847,546 | 64,673,114 |
| Average % of the lane | 0.64% | 0.53% | 37.68% | 42.14% |
| Percent perfect index reads | 96.99% | 94.13% | 91.72% | 92.64% |
| Mean transcript Quality Score | 35.57 | 35.53 | 33.06 | 33.12 |
| Mean Barcodes and UMI quality score | 36.22 | 36.19 | 33.02 | 33.14 |

### TruDrop libraries maintain high quality when multiplexed in a high throughput fashion

With the successful testing of the two initial pairs of indices on the NovaSeq, 24 human and mouse samples were prepared and sequenced, each uniquely dual-indexed, on the NovaSeq 6000 alongside 186 other Illumina libraries. TruDrop libraries yielded 94%-151% of the target 125 million reads per sample (Supplementary Table 3). In total, the 24 samples represented 29.4% of the raw sequencing yield across all of the lanes from the flow cell, equivalent to sequencing alongside ~70% PhiX on the NextSeq. Based on our prior sequencing results of the V2 libraries alongside PhiX on the NextSeq, we would therefore have expected to see a decrease in the read quality scores compared with the 2-sample run due to a decrease in the of libraries represented on the flow cell. However, the quality scores and error rates were observed to be very similar. The average transcript and barcodes + UMI quality scores were 35.32 and 36.07, respectively, (Supplementary Table 3). These do not differ greatly from the prior TruDrop NovaSeq sequencing run (Table 2) and are still a 2.0- and 1.7-fold reduction in base call error rate over V2 libraries on the NextSeq with PhiX, and a 3.6- and 2.9-fold reduction in error over just V2 libraries alone on the NextSeq. These results suggest that the improved quality scores observed on the NovaSeq can be maintained as long as some minimum diversity of Illumina libraries are present. The base calling accuracy plot also confirms this improvement in base calling accuracy, as the region covering the cell barcodes + UMI (cycles 1-11 and 31-50) displays more than 90% of the reads were above Q30 (Fig. 4C). For the first 100 transcript read bases, 90% or more of the reads were at or above Q30. The drop observed in the base calling accuracy plot at cycle 60 that continues to the end of read 1 (cycle 150), corresponding with where the poly T capture sequence is located. This decrease in accuracy only continued through regions that would be trimmed out during mapping and barcode deconvolution. The decrease in accuracy did not affect other Illumina libraries on the flow cell, as when considered individually, 95% of other Illumina libraries had greater than 90% of reads at or above Q30 for the entire sequencing run. These results demonstrate that up to 24 TruDrop libraries can be multiplexed on the NovaSeq alongside standard Illumina libraries, while maintaining a very high sequencing quality for both inDrop and Illumina libraries. With lane splitting, 4 pools of 24 samples can be sequenced across 4 sequencing lanes for a total of 96 inDrop libraries sequenced at a time.

### TruDrop libraries sees sequence alignment rates

To investigate if the improvement in base call accuracy had a measurable effect on downstream data quality, two colonic (one mouse and one human) libraries that had previously been sequenced as V2 libraries on the NextSeq were re-made with the TruDrop structure and sequenced on the NovaSeq. The reads for the sequenced V2 libraries and the TruDrop libraries were then aligned and deconvolved in parallel. The overall percentage of reads that aligned did not significantly change from V2 to TruDrop libraries for either the mouse (96.38% and 96.56%, respectively respectively) or human (96.11% and 95.12%, respectively) replicates (Table 3). However, for the mouse sample the percentage of unique alignments increased from 67.15% to 73.48%, while the human sample experienced a similar improvement from 84.44% to 87.23%. The improved rates of uniquely aligned reads have been consistent to date for all TruDrop samples sequenced on the NovaSeq (data not shown).

**Table 3.** Comparison of data alignment quality of the V2 and TruDrop structures

| Sample | Sequencer | Sequencing Depth (reads) | mapped reads (%) | Uniquely aligned reads (%) |
| --- | --- | --- | --- | --- |
| V2 Mouse | NextSeq | 98606967 | 96.38 | 67.15 |
| TruDrop Mouse | NovaSeq | 43657381 | 96.56 | 73.48 |
| V2 Human | NextSeq | 55507773 | 96.11 | 84.44 |
| TruDrop Human | NovaSeq | 188061057 | 95.12 | 87.23 |

### Single-cell data generated by TruDrop maintain the same cell population structure as V2

To determine whether scRNA-seq data generated with TruDrop was valid, count data were generated by alignment, deconvolution, and filtering in a manner parallel to the same samples generated with V2. For sets of mouse and human samples, data generated by the two library structures were analyzed together using t-SNE (22) to reveal significant mixing between TruDrop and V2, with identical cell types detected (Fig. 5A, C, Supplementary Fig. 2A). To quantify this mixing, we used sc-UniFrac (23), a distance metric between 0 and 1, with 0 signifying two samples to be identical and 1 signifying complete non-overlap. For all sets samples (mouse/human), the sc-UniFrac distance is 0.07, strongly suggesting that cell populations identified with the different libraries are almost completely identical (Fig. 5B, D Supplementary Fig. 2B), with minor differences (such as in erythrocytes) due to the small number of cells in those clusters. These data suggest that the library structure and sequencer used did not result in any overt biases in data for recovering cell types.

**Figure 5.**
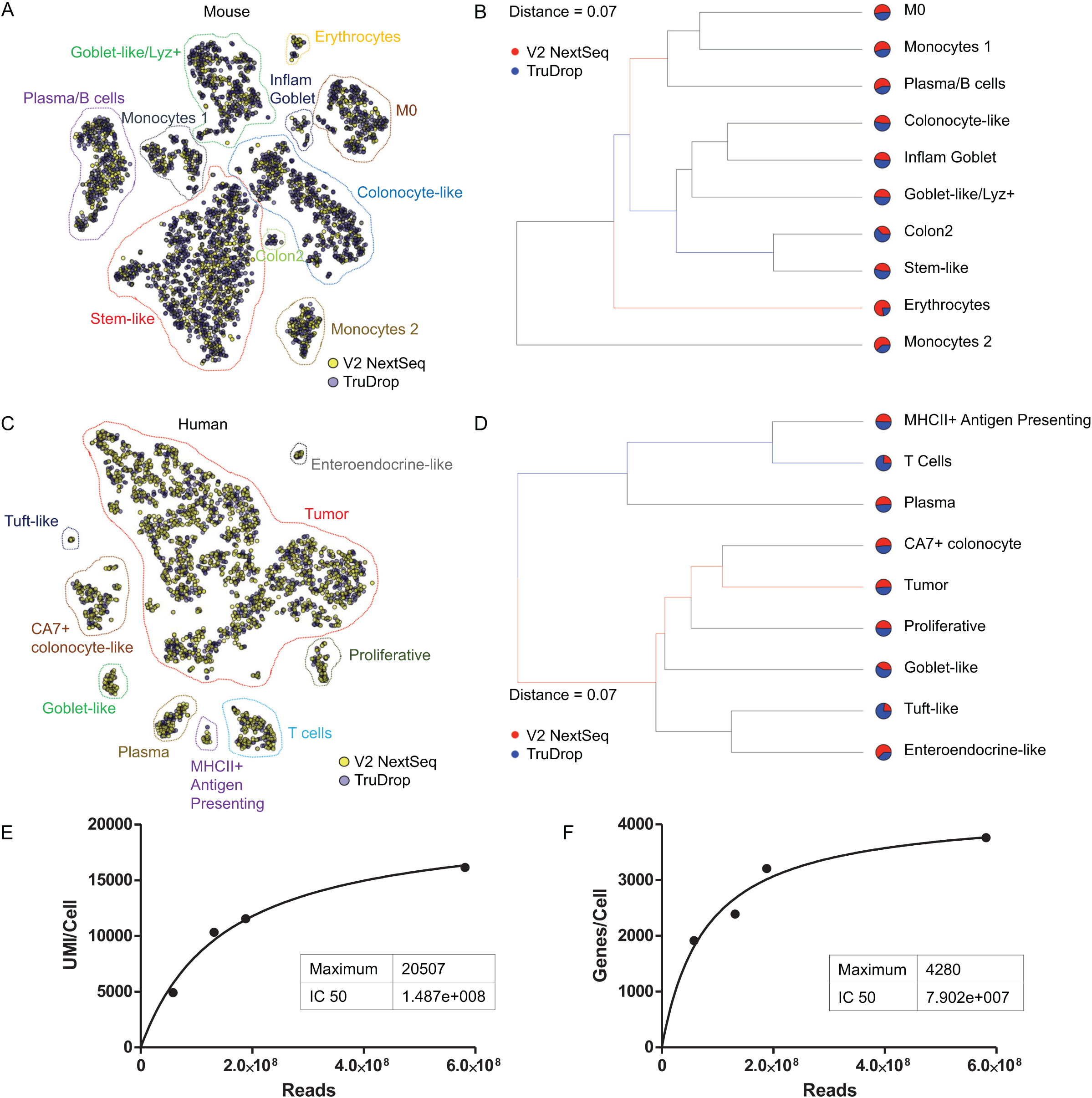
Comparison of cell types identified between V2 libraries on NextSeq and TruDrop on NovaSeq. (A and C) Combined t-SNE analysis of cells identified from a TruDrop and V2 library prepared from the same samples of (A) mouse and (C) human tumors. (B and D) sc-UniFrac tree representations of subpopulation structures for libraries presented in A and C, respectively. Cell groups enriched using V2 NextSeq libraries have red branches, while those enriched using TruDrop NovaSeq have blue branches. Thickness of branches represent level of enrichment. Distance values range from 0 to 1, with 0 representing complete overlap between two datasets. (E) Median UMI/cell and (F) genes/cell detected as a function of read depth using TruDrop on the NovaSeq. The maximum UMI/Cell and genes/cell are predicted by hyperbolic curve fitting.

### TruDrop libraries generate larger throughput of data on the NovaSeq

We evaluated the performance of TruDrop libraries of human colonic specimens at different sequencing depths by comparing the number of UMIs and genes recovered after NovaSeq sequencing (Fig. 5E, F). Similar to prior testing, diminishing returns were observed with increasing read depth due to re-sequencing of reads that collapse into single UMIs (3). In this prior work, medians of ~3,000 UMI/cell and ~1,300 genes/cell were reported when samples were sequenced to ~60K reads per cell, with a predicted maximum of ~3,500 UMI/cell and ~1,400 genes/cell (3). For the samples sequenced here, we observed medians of ~16,000 UMI/cell and ~3,800 genes/cell when samples were sequenced to 150K reads per cell (Supplementary Table 4). The predicted maximum output in our runs is 20,507 UMI/cell and 4,280 genes/cell (Fig. 5E, F). While cell typing could be done with as few as ~20K reads per cell (Fig. 5A–D), we find that analysis in the range of 40K to 60K reads per cell (~11,000 UMI/cell, ~2800 genes/cell) yields the most return for value.

## Discussion

Multiplexed NGS is currently essential for performing scRNA-seq in a cost-efficient manner. In order fully realize the advantage of the decreased costs associated with sequencing on platforms that utilize Illumina’s ExAMP chemistry, it is necessary for scRNA-seq libraries to utilize a multiplex sequencing strategy that adequately addresses the problem of index hopping. With the development of TruDrop, we take a preventative approach in utilizing a unique dual-indexing method that minimizes sample cross-talk (6). Most prior work on high-throughput scRNA-seq libraries has focused on using computational methods to deconvolve and filter out entire barcodes (cells) with reads that could have originated from index-hopped sequencing reads, resulting in substantial data loss (9). To our knowledge only the V3 inDrop library structure has previously endeavored to implement a dual-indexed system for high-throughput scRNA-seq (8). Its use of a portion of the cell barcode as the i7 index, however, means that the i7 index could be repeated across samples. It was thus a combinatorial dual-indexed system that would not resolve the cross-sample barcode collision problem. The work documented here allows for the independent evaluation of samples when filtering for barcode collisions, resulting in an increased retention of cell barcodes compared with that of single-indexed samples. Users who do not have access to the NovaSeq can also use this dual-indexed design for decreased cross-sample contamination on the HiSeq 3000, HiSeq 4000, and HiSeq X Ten, which also rely on patterned flow cells and ExAmp chemistry. Meanwhile, users who are restricted to sequencing inDrop libraries on the NextSeq platform, but still wish to use standard Illumina sequencing primers can use a single-indexed version via the universal TruSeq P5 (cell barcode + UMI) structure.

Sequencing inDrop libraries alongside libraries with a diverse base composition on the NovaSeq results in much lower (3.7-fold decrease) base-calling error rates compared with those observed on the NextSeq. This substantial improvement of sequencing quality is maintained when 24 TruDrop samples (30% of a run) were sequenced alongside Illumina libraries, with no effect on the quality of the standard libraries. The reduction in the base-calling error rate observed with the TruDrop on the NovaSeq is likely the major contributor to the increase in percent of uniquely aligned reads to the reference genome, as more accurate reads should result in a lower rate of ambiguous alignments. The uniquely aligned reads are those that move on to downstream data analysis, and thus, this improvement results in substantially more useable data. As for the discrepancy in the percentage of uniquely aligned reads between mouse (73%) and human (87%), this is a routinely observed difference between mapping to reference genomes of mouse versus human. Furthermore, the TruDrop libraries did not generate biased results, as sequencing the same samples using either library structures recovered the same cell types, with TruDrop libraries producing higher quality data.

In summary, the TruDrop library structure resulted in the ability to sequence inDrop libraries on the NovaSeq by solving the problem of index hopping. The resulting sequencing data have lower base call error rates, likely due to increased diversity of libraries sequenced from high multiplexity, resulting in better sequence alignments. The adoption of high-throughput next generation sequencing technologies results in substantial cost savings that should enable large scale cohort studies, with hundreds of samples, to be assayed by scRNA-seq.

## Conflict of Interest Statement

Y.Z., J.X., E.B.P., X.C. are employees of RootPath, Inc. M.J.B, C.J.H.B, L.N.M.Q. are employees of 1CellBio, Inc. Individuals listed above played no role in comparative analyses performed in this paper.

## Acknowledgements

K.S.L is funded by R01DK103831 and U01CA215798. A.N.S. and A.J.S. are funded by U2CCA233291 and U54CA217450. Q.L., K.S.L., and M.A.R.S. are funded by P50CA236733. P.N.V. is supported by T32HD007502. B.C. is supported by T32LM012412. C.R.S. was supported by T32AI007281. We thank Linas Mazutis from MSKCC for his valuable input on the library preparation protocol, Karen Beeri from the VANTAGE core for her technical assistance, and members of the Lau lab, Vanderbilt Epithelial Biology Center, and Quantitative Systems Biology Center for helpful discussions.

---

**Supplementary Figure 1.**
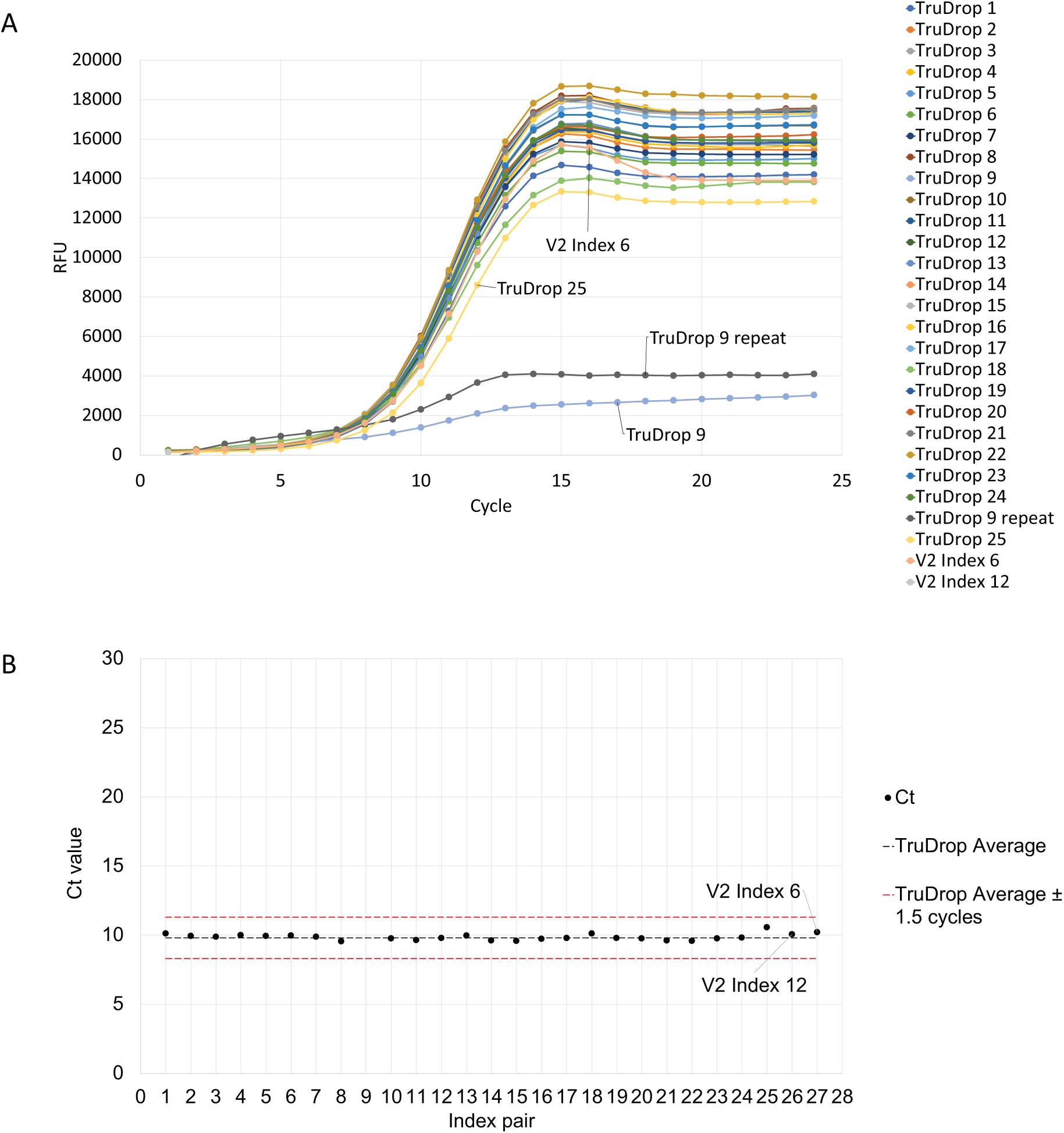
Comparison of amplification of TruDrop and V2 primers during library preparation. (A) Diagnostic qPCR amplification curves comparing performance of all TruDrop primer pairs to V2 primers, all performed on the same sample. (B) Ct values of A.

**Supplementary Figure 2.**
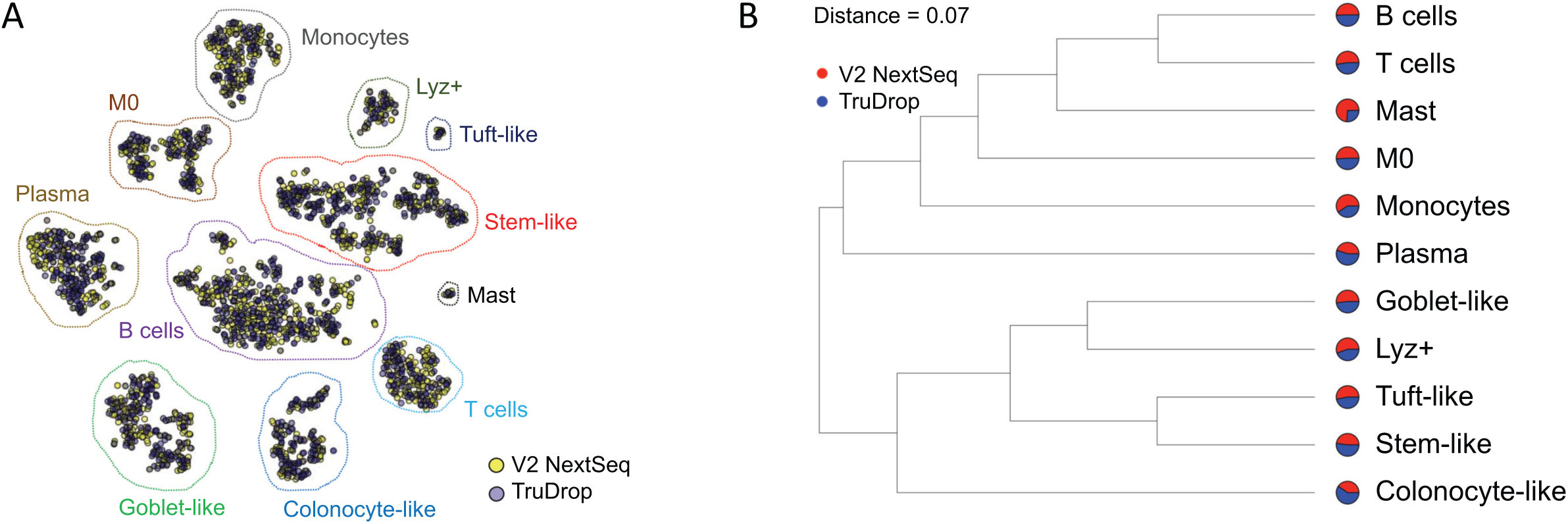
Another example comparison of cell types identified between V2 on NextSeq and TruDrop on NovaSeq. (A) t-SNE and (B) sc-UniFrac analysis as performed in Figure 5.

**Supplementary Table 1.** Cost of Sequencing for inDrop^†^

| Sequencer | Flow cell | Sequencing Kit | Cost of Flow cell | Number of lanes | Possible Number of Samples per flow cell | Sequencing Cost per sample |
| --- | --- | --- | --- | --- | --- | --- |
| NextSeq | High Throughput | PE 75 | \$3055.00 | 4 | 4 | \$764.75 |
| NovaSeq | S2** | PE 150 | \$9,840.00** | 2 | 37 | \$531.89 |
| NovaSeq | S4 | PE 150 | \$36,135.00 | 4 | 96* | \$361.35 |

<sup>†</sup>assuming a read dept of 100 million reads per sample and unless otherwise noted costs are from local sequencing core facility
\*Assumes sharing flow cell with other users
\*\* requires lane splitting and cost is pulled from University of Wisconsin
(https://www.biotech.wisc.edu/services/dnaseq/pricing)

**Supplementary Table 2.** Evaluation of two TruDrop libraries’ raw yield and quality in low-throughput sequencing run on the iSeq 100

| Library | Sequencer | i7 | I5 | Expected Reads | Observed inDrop Reads* |
| --- | --- | --- | --- | --- | --- |
| Mouse 4 | iSeq 100 | CCGCGGTT | AGCGCTAG | 2,000,000 | 2,876,464 |
| Mouse 5 | iSeq 100 | TTATAACC | GATATCGA | 2,000,000 | 3,166,938 |

\*Libraries were sequenced alongside a 10% spike-in of PhiX.
The TruSeq-inDrop (TruDrop) libraries using both an i7 and i5 index saw about 1.5x the expected yield on the iSeq sequencer indicating that for this test the flow cell over-clustered. The iSeq uses similar chemistry to that of the NovaSeq. This shows that the TruDrop structured libraries could be sequenced on the NovaSeq.

**Supplementary Table 3.** 24 TruDrop libraries raw data yield and quality in combined high-throughput sequencing run on the NovaSeq

| Sample | i7 | i5 | Expected<br>inDrop Reads | Observed<br>inDrop<br>reads | Average<br>% of the<br>lane | perfect<br>index<br>read (%) | Mean Barcodes<br>and UMI<br>Quality Score | Mean<br>Transcript<br>Quality Scores |
| --- | --- | --- | --- | --- | --- | --- | --- | --- |
| 1 | CCGCGGTT | AGCGCTAG | 125,000,000 | 163,510,352 | 1.38 | 96.84 | 36.05 | 35.36 |
| 2 | TTATAACC | GATATCGA | 125,000,000 | 141,764,120 | 1.20 | 96.44 | 36.02 | 35.34 |
| 3 | GGACTTGG | CGCAGACG | 125,000,000 | 127,912,777 | 1.08 | 96.91 | 35.97 | 35.44 |
| 4 | AAGTCCAA | TATGAGTA | 125,000,000 | 141,900,719 | 1.20 | 96.83 | 36.03 | 35.41 |
| 5 | ATCCACTG | AGGTGCGT | 125,000,000 | 153,271,668 | 1.29 | 97.04 | 36.02 | 35.36 |
| 6 | GCTTGTCA | GAACATAC | 125,000,000 | 131,683,586 | 1.11 | 96.71 | 36.09 | 35.16 |
| 7 | CAAGCTAG | ACATAGCG | 125,000,000 | 168,538,426 | 1.42 | 96.84 | 36.10 | 35.43 |
| 8 | TGGATCGA | GTGCGATA | 125,000,000 | 124,903,031 | 1.05 | 97.61 | 36.04 | 35.42 |
| 9 | GACCTGAA | TTGGTGAG | 125,000,000 | 125,928,312 | 1.08 | 95.63 | 36.07 | 35.24 |
| 10 | TCTCTACT | CGCGGTTC | 125,000,000 | 132,604,827 | 1.13 | 96.32 | 36.10 | 35.31 |
| 11 | CTCTCGTC | TATAACCT | 125,000,000 | 121,444,989 | 1.03 | 96.79 | 36.16 | 35.53 |
| 12 | CCAAGTCT | AAGGATGA | 125,000,000 | 127,434,526 | 1.08 | 96.80 | 36.12 | 35.48 |
| 13 | TTGGACTC | GGAAGCAG | 125,000,000 | 143,214,632 | 1.21 | 97.09 | 36.08 | 35.46 |
| 14 | GGCTTAAG | TCGTGACC | 125,000,000 | 121,001,482 | 1.03 | 96.30 | 36.09 | 35.34 |
| 15 | AATCCGGA | CTACAGTT | 125,000,000 | 117,718,028 | 1.00 | 96.35 | 36.10 | 34.59 |
| 16 | TAATACAG | ATATTCAC | 125,000,000 | 176,705,278 | 1.50 | 96.49 | 36.01 | 35.30 |
| 17 | CGGCGTGA | GCGCCTGT | 125,000,000 | 176,054,943 | 1.51 | 95.49 | 36.04 | 35.37 |
| 18 | ATGTAAGT | ACTCTATG | 125,000,000 | 164,005,038 | 1.38 | 97.05 | 36.11 | 35.20 |
| 19 | GCACGGAC | GTCTCGCA | 125,000,000 | 150,680,775 | 1.26 | 97.57 | 36.04 | 35.31 |
| 20 | GGTACCTT | AAGACGTC | 125,000,000 | 170,171,924 | 1.43 | 97.14 | 36.09 | 35.29 |
| 21 | AACGTTCC | GGAGTACT | 125,000,000 | 128,785,547 | 1.09 | 96.74 | 36.12 | 35.36 |
| 22 | GCAGAATT | ACCGGCCA | 125,000,000 | 188,900,350 | 1.61 | 95.62 | 36.11 | 35.45 |
| 23 | ATGAGGCC | GTTAATTG | 125,000,000 | 126,939,171 | 1.08 | 96.08 | 36.06 | 35.30 |
| 24 | ACTAAGAT | AACCGCGG | 125,000,000 | 150,500,764 | 1.31 | 93.28 | 36.14 | 35.18 |

**Supplementary Table 4.** Diversity of UMI’s and genes expressed for cells sequenced with the TruDrop structure

| Sample | Reads/cell<br>encapsulated | Cells<br>detected | Median UMI's/Cell<br>(25<sup>th</sup> percentile, 75<sup>th</sup><br>percentile) | Median Genes/Cell<br>(25<sup>th</sup> percentile, 75<sup>th</sup><br>percentile) |
| --- | --- | --- | --- | --- |
| TruDrop Human 1 | 19398 | 1246 | 4902 (3162, 7074) | 1916 (1292, 2498) |
| TruDrop Human 2 | 39302 | 927 | 10332 (6178, 15303) | 2393 (1674, 3241) |
| TruDrop Human 3 | 61974 | 1381 | 11545 (8043, 15921) | 3208 (2382, 4207) |
| TruDrop Human 4 | 149535 | 2049 | 16141 (12034, 21397) | 3760 (2985, 4537) |

## Supplementary text

### Rationale of library and primer design

The standard Illumina TruSeq library incorporates the following adapter sequences on either end of the library respectively: P7: 5’ – CAAGCAGAAGACGGCATACGAGAT[i7]GTGACTGGAGTTCAGACGTGTGCTCTTCCGATCT – 3’ P5: 5’ – AATGATACGGCGACCACCGAGATCTACAC[i5]ACACTCTTTCCCTACACGACGCTCTTCCGATCT – 3’.

The sequence present on the 5’ side of the i7 and i5 indexes are the adapter sequence required for annealing and cluster formation on the Illumina flow cell. The sequences to the 3’ side of the i7 and i5 indexes are where the TruSeq sequencing primers will bind during the sequencing process.

The sequence of the V2 inDrop library structure is as follows: Cell Barcode + UMI(P7): 5’ – CAAGCAGAAGACGGCATACGAGAT [i7] CTCTTTCCCTACACGACGCTCTTCCGATCT [cell barcode 1] GAGTGATTGCTTGTGACGCCTT [Cell barcode 2] [UMI] TTTTTTTTTTTTTTTTTTT… – 3’ Transcript (P5): 5’ – AATGATACGGCGACCACCGAGATCTACACGGTCTCGGCATTCCTGCTGAACCGCTCTTCCGATCTNNNN NN… – 3’.

For the cell barcode + UMI side of the V2 library structure, a truncated version of the Illumina i5 sequencing primer site was used as the sequencing primer for the cell barcode + UMI (P7 side). On the P5 - transcript side of the V2 inDrop library, a sequencing primer site that is currently considered obsolete by Illumina was used. This obsolete priming site on the P5 side of the V2 structure is added on via the use of a random hexamer during the 2^nd^ RT and is then extended to the complete P5 V2 structure during a brief PCR. The truncated P5 sequencing priming site used on the P7 side of the V2 library is partly built into the primer sequence attached to the hydrogel bead used to capture the transcriptomic material during encapsulation. This truncated Illumina P5 primer sequence used on the P7 side has 12 bases in common with the full length standard Illumina P7 primer sequenced. This will likely result in mis-priming events on inDrop libraries when sequencing V2 inDrop alongside large numbers of Illumina libraries. The P5 side of the V2 structure could be changed due to its priming with a random hexamer. The P7 side could be changed so long as the resulting structure used the Illumina P5 sequencing primer site present on the primer used by the V2 hydrogel beads.

For the new TruSeq-inDrop (TruDrop) library structure the P7 and P5 sides were swapped so that the sequencing primer and flow cell binding site for the cell barcode + UMI side of the library followed Illumina’s TruSeq libraries. The transcript side of the library now uses the P7 structure of TruSeq. The sequence for the final TruDrop library is as follows: Transcript (P7): 5’ – CAAGCAGAAGACGGCATACGAGAT [i7] GTGACTGGAGTTCAGACGTGTGCTCTTCCGATCTNNNNNN… – 3’ Cell Barcode + UMI (P5): 5’ – AATGATACGGCGACCACCGAGATCTACAC [i5] ACACTCTTTCCCTACACGACGCTCTTCCGATCT [cell barcode 1] GAGTGATTGCTTGTGACGCCTT [cell barcode 2] [UMI] TTTTTTTTTTTTTTTTTTT… −3’.

The new TruDrop library structure utilizes the standard Illumina TruSeq sequencing primers. It also incorporates a unique i7 and unique i5 index for each sample. The i7 and i5 index pairs were picked from the set of 96 pairs of unique dual indexes that Illumina has published as the “IDT for Illumina TruSeq UD Indexes”. The TruDrop library preparation follows the same steps as previously published for the V2 library with the substitution of the following primers for their V2 counterparts: TruDrop 2^nd^ RT primer: 5’ – GTGACTGGAGTTCAGACGTGTGCTCTTCCGATCTNNNNNN – 3’ TruDrop PE1: 5’ – AATGATACGGCGACCACCGAGATCTACAC [i5] ACACTCTTTCCCTACACGA – 3’ TruDrop PE2: 5’ – CAAGCAGAAGACGGCATACGAGAT [i7] GTGACTGGAGTTCAGACGTGT – 3’. TruDrop 2^nd^ RT primer was ordered from IDT as desalted. TruDrop PE1 and PE2 primers were all ordered from IDT as TruGrade HPLC purified primers in individual tubes to minimize risk of cross-contamination during synthesis and handling. V2 PE2-N6 primer was ordered as desalted from Sigma. V2 PE1 and PE2 primers were ordered PAGE purified from Sigma. Primers were all resuspended at 100 µM in 10 mM Tris-HCl pH 8.0 and 0.1 mM EDTA pH 8.0. PE1 and PE2 primers were then diluted to 10 µM. For V2 libraries PE1 was mixed with PE2 in a 1:1 ratio (concentration of 5 µM for each primer) for working aliquots. For TruDrop libraries, unique dual-index primer pairs were then mixed in 1:1 ratio (concentration of 5 µM for each primer) for working aliquots.

## Methods

### Calculation of cross-sample barcode collision as a result of index hopping

An estimate of the number of barcodes/cells to be thrown out per sample can be calculated as follows. A prior study (1) documents the index hopping rate on a NovaSeq run to be 4.85%. Assuming it is equally likely for any given read to hop from one sample to the next, all of the samples should be treated as if all of the cells that they contain belong to a single sample. The manner of calculating rates of barcode collision for inDrop libraries was previously documented by (2–5). Rates of barcode collision for pools of 2, 4, 12, 24, and 48 samples (6000, 120000, 36000, 72000, and 144000 cells respectively). Barcode collision and index hopping are 2 independent events so the probability of either occurring in a set number of cells is *P(barcode collision) + P(index hope) − P(barcode collision and index hop)*. The resulting rate represents the percentage of cell barcodes discarded due to cross-sample barcode collision.

### Mouse Colonic Crypt Isolation and Dissociation

All animal protocols were approved by the Vanderbilt University Animal Care and Use Committee and in accordance with NIH guidelines. *Lrig1*^*CreERT2*^ and *Apc*^*fl*^ mice on C57BL/6 background were purchased from Jackson Laboratory. At 12 weeks mice received 1-3 colonoscopy guided orthotropic injections of 0.70 mL of 100µM 4-hydroxytamoxifen. The following day mice were administered 2.5% DSS (TdB consultancy, batch DB001-37) in deionized water for 6 days in their drinking water. Mice were sacrificed 28 days following 4-hydroxytamoxifen injections. Colonic tumors were dissected and incubated in chelation buffer (3mM EDTA, 0.5 mM DTT) at 4°C for 1 hour 15 minutes. The tissue was shaken in 10 mL of PBS in a 15 mL conical tube for 2 minutes to release the crypts. The crypt suspension was centrifuged at 250-300 xg for 5 min at 4°C. Crypts were washed three times with 1x DPBS. The crypts were dissociated into single cells using a cold-activated protease (1 mg/mL) and DNase I (2.5 mg/mL) mixture in 1x DPBS on a rocker at 4°C. The cells were then washed three times with 1x DPBS after spinning 600x g for 5 min each at 4°C.

### Human Colonic Crypt Isolation and Dissociation

All studies were performed according to Vanderbilt University Institutional Review Board. Colonic biopsies were collected and placed into RPMI or UWA prior to processing. Upon arrival biopsies were minced to 4 mm^2^ and washed with 1x DPBS. They were then incubated in chelation buffer (4mM EDTA, 0.5 mM DTT) at 4°C for 1 hour 15 minutes. The tissue was then dissociated with cold protease and DNase I for 25 min. Single-cell suspensions were triturated at the start and every 10 minutes with a P1000 pipette tip with the tip 0.1-0.5 cm removed. Single cells were washed three times with 1x DPBS after spinning 600X g for 5 min each at 4°C.

### inDrop Single-Cell Encapsulation and Library Preparation

A target of 3000 single cells per sample were encapsulated and barcoded using the inDrop platform with 1Cell-Bio library preparation protocol version 2.3. Modifications to the protocol include reverse transcription as noted in (6), ExoI digestion, second strand synthesis, and T7 *in vitro* transcription as noted in version 1.2. Furthermore, we doubled the volumes of diagnostic qPCR and final PCR steps, with a final double-sized size selection. For TruDrop-specific modifications, we used TruDrop custom primers (RT, PE1, PE2).

### TruDrop Primer Testing via qPCR

To test if the efficiency of TruDrop dual indexing primers, a single mouse inDrop library was prepared up through the second RT using the TruDrop RT primer. The sample was used to run a diagnostic qPCR each pair of TruDrop i7 and i5 indexes, all in parallel, on a BioRad C1000 Touch Thermal Cycler CFX96 Real-time system. To verify that the TruDrop primers amplified appropriately, we compared their amplification curves with two V2 libraries that had previously produced good results on the NextSeq. An index pair not reaching the Ct value of 5000 RFU was not included in subsequent analysis. Based off of prior testing by (7), it was expected that the Ct for individual primer pairs would not deviate from the average by more than 1.5 cycles.

### Illumina Sequencing

All libraries were evaluated on a Qubit 3.0 fluorometer and an Agilent 2100 Bioanalyzer regarding concentration and fragment size distribution prior to sequencing on various platforms.

*NextSeq:* V2 libraries were sequenced on the NextSeq 500 using a PE 75 kit in a customized sequencing run as previous (Herring et al., 2018). 10-15% PhiX was pooled when appropriate. *MiSeq:* Sequencing of a V2 library on the MiSeq was performed using the Reagent Kit v2 Nano with custom sequencing primers, along with a 10% PhiX spike-in. Sequencing was performed using 30 cycles for read 1 (transcript), 6 cycles for the index read, and 30 cycles for read 2 (cell barcode + UMI).

*iSeq 100:* TruDrop libraries were sequenced on the iSeq with a 10% PhiX spike-in using a PE 150 kit. The cell barcode + UMI was sequenced on read 1. The transcript was sequenced on read 2. *NovaSeq 6000:* Sequencing on the NovaSeq was performed using a S4 flow cell with a PE 150 kit. TruDrop libraries, at a 2nM standard loading concentration, were pooled with other Illumina compatible libraries, and sequenced to various target depths (50 – 500 million reads).

### Downstream data analysis

For all sequence data, reads were demultiplexed using bcl2fastq v2.20.0.422. Base call accuracy (% >= Q30 score) plots were generated via Illumina’ BaseSpace. Quality scores were generated using fastQC to find the average quality score per cycle for reads from the demultiplexed fastq files (8). The proportion for how much each cycle was contributing to each transcript, barcode 1, barcode 2, and UMI read was determined and used to calculated the weighted average of the quality score for the transcript (first 100 bases only) and cell barcodes + UMI. Base call error rates were then calculated using the formula *p* = 10^(−*Q*/10)^.

Following demultiplexing, reads were filtered, sorted by their barcode of origin, and aligned to the reference transcriptome to generate a counts matrix using the DropEst pipeline (9). Barcodes containing cells were filtered for further analysis, as previous (10), and aligned using Harmony (11). t-SNE and sc-UniFrac analyses were performed following previous methods (10, 12) in Matlab (Mathworks) and R, respectively.

